# Proteomic analysis of the actin cortex in interphase and mitosis

**DOI:** 10.1101/2022.02.16.480677

**Authors:** Neža Vadnjal, Sami Nourreddine, Geneviève Lavoie, Murielle Serres, Philippe P. Roux, Ewa K. Paluch

## Abstract

In animal cells, many shape changes are driven by gradients in the contractile tension of the actomyosin cortex, a thin cytoskeletal network that supports the plasma membrane. Elucidating how cortical tension is controlled is thus essential for understanding cell and tissue morphogenesis. Increasing evidence shows that alongside myosin II activity, actin network organisation and composition are key to cortex tension regulation. However, how cortex composition changes when cortical tension changes remains poorly understood. Here, we compared cortices from cells in interphase and in mitosis, as mitosis entry is associated with a strong increase in cortical tension. We purified cortex-enriched cellular fractions and analysed their composition using mass spectrometry, identifying 922 proteins consistently represented in both interphase and mitotic cortices. We curated this dataset by focusing on actin-related proteins, narrowing down to 238 candidate regulators of the cortex during the mitotic cortical tension increase. Among these candidates, our analysis pointed to a role for septins, and in particular septin 9, in the control of mitotic cell rounding. Overall, our study brings insight into the regulation of mitotic rounding, and paves the way for systematic investigations of the regulation of cell surface mechanics.

## Introduction

Changes in animal cell shape are largely controlled by the actin cytoskeleton. In particular, the actomyosin cortex, a thin (150 nm thick) network of actin, myosin and associated proteins underlying the plasma membrane, drives contraction-based cellular deformations (1, 2). Contractile tension is generated in the cortex by myosin motors, pulling on actin filaments, and increasing cortical tension promotes cell rounding. For instance, a strong increase in cortical tension at the onset of mitosis typically leads cells to round up in prometaphase (3, 4). The resulting mitotic rounded shape provides the space required for spindle assembly, positioning, and accurate chromosome separation (5). Furthermore, gradients in cortical tension lead to local contractions and induce cell shape changes driving processes such as cell division, cell migration, and epithelial morphogenesis (2, 6). Given its importance in the regulation of cell shape, it is thus essential to systematically identify the key factors controlling cortical tension.

The role of myosin motors in tension generation has been extensively characterised (7–10). However, a number of recent studies have highlighted the importance of other actin-binding proteins. In particular, regulators of actin filament nucleation and length, as well as crosslinkers have been shown to play a key role in the control of cortical tension (11– 14). Furthermore, intermediate filaments have been shown to contribute to the cortical tension increase observed in mitotic cells (15, 16). Taken together, the emerging view is that cortical tension is controlled by a variety of mechanisms that modulate myosin activity, actin network organisation, as well as cortex interactions with other cytoskeletal networks. However, systematic investigations of cortex tension regulation have been hindered by our limited understanding of how cortical composition changes when tension changes.

Here, we set out to systematically investigate which actin-related proteins display different levels at the cortex in cells with high and low tension. To this aim, we analysed the proteic composition of actin cortices in cells synchronised in interphase and mitosis, as mitotic cells display considerably higher cortical tension compared to interphase cells (12, 17). We used purified cellular blebs, as we have previously shown that blebs assemble a cortex similar to the cellular cortex, and can be isolated from cells, yielding cortex-enriched cellular fractions (18, 19). We compared, using mass spectrometry, the composition of blebs separated from interphase and mitotic cells. We identified 922 different proteins consistently present in blebs from both mitotic and interphase cells, among which 238 were actin-related proteins. Finally, we showed that one of the candidates identified, septin 9, regulates mitotic cell rounding. Taken together, our systematic analysis generates a list of candidate regulators of the cortical remodelling taking place at mitotic entry. It also identifies septins, and in particular septin 9, as important regulators of cortex-driven cell rounding at the transition between interphase and mitosis.

## Results

### Isolation of actin cortex-enriched blebs from cells synchronised in interphase and mitosis

To obtain interphase cells, HeLa cells were synchronised in G1/S phase using thymidine (Figure 1A, see Methods for details). Synchronised interphase cells were then detached and maintained in suspension (Figure 1B, upper panel), as under these conditions interphase cells display a continuous cortex and a cortical tension considerably lower than in mitotic cells (12). To obtain mitotic cells, HeLa cells were synchronised in prometaphase using S-Trityl-L-Cystine (STLC). Mitotic cells were further enriched by mechanically separating them from adherent cells through a mitotic shake-off, leading to a population of rounded mitotic cells (Figure 1B, bottom panel). We checked synchronisation efficiency by DNA staining of the synchronised cell populations (Figure 1B), and by immunoblotting for levels of the mitotic markers phosphorylated histone H3 and cyclin B (Figure 1C). This confirmed that our synchronisation protocol yielded cell populations of predominantly rounded interphase and mitotic cells.

**Fig. 1.**
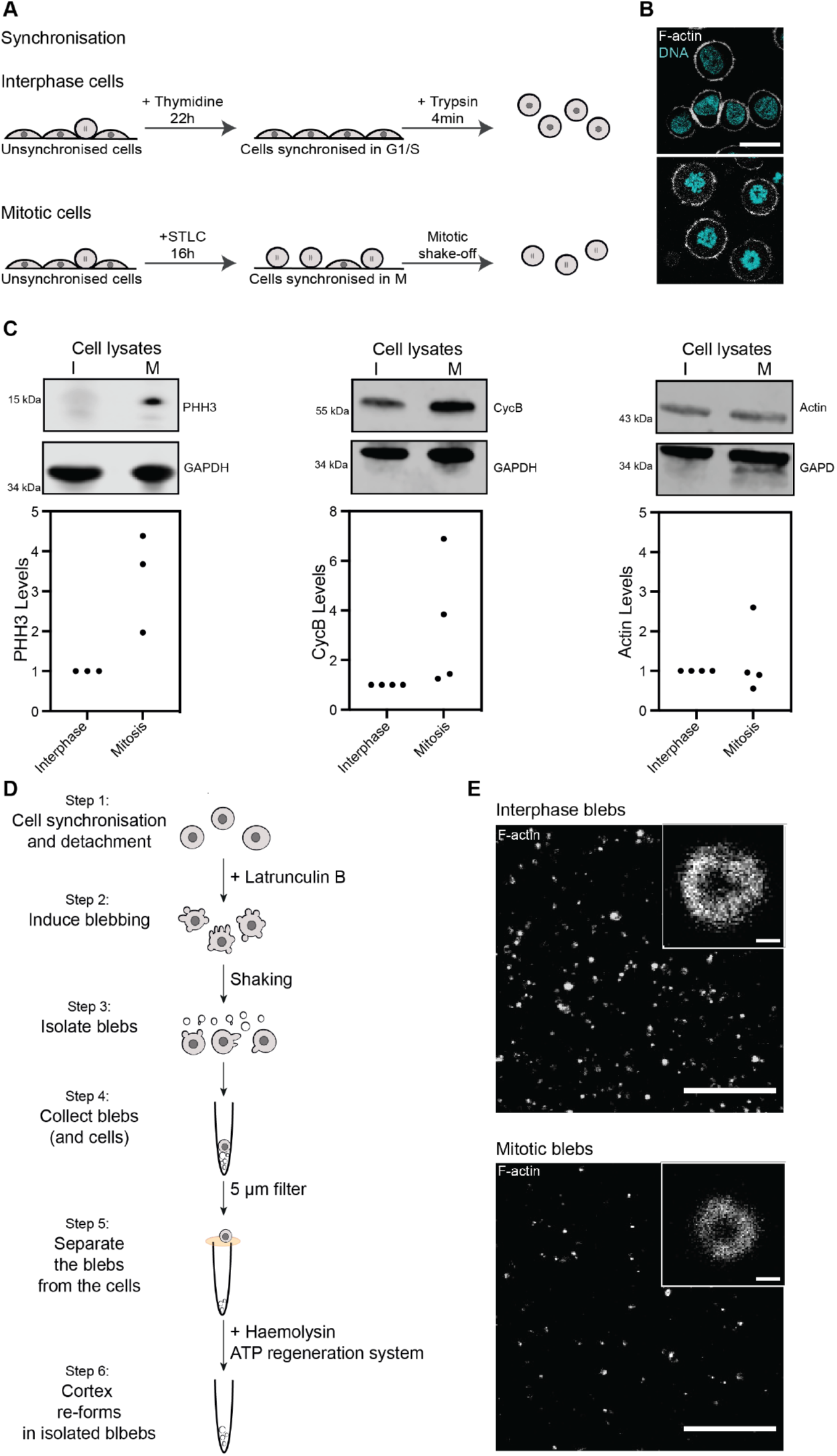
Isolation of cortex-enriched blebs from interphase and mitotic cells. (A) Schematic describing cell synchronisation in interphase (top) and mitosis (bottom). Prior to bleb isolation, cells were synchronised in interphase (G1/S phase) with a 22 h thymidine treatment, detached from the dish and rounded up using trypsin. For synchronisation in mitosis, the cell population was first enriched for mitotic cells with a 16 h treatment with the Eg5 inhibitor STLC, which prevents bipolar spindle formation, followed by a mitotic shake-off. (B) Representative confocal images of synchronised interphase (upper panel) and mitotic (lower panel) cells. Cyan: DAPI (DNA), white: Phalloidin (F-actin). DAPI staining shows the nuclear organisation of DNA in cells synchronised in interphase and condensed chromosomes in cells synchronised in mitosis. Scale bar = 20 µm. (C) Western blot analysis of interphase and mitotic cell lysates for the mitotic markers phosphorylated histone H3 and cyclin B, and for actin, confirming synchronisation at the cell population level. GAPDH: loading control. (D) Schematic depicting the bleb isolation protocol. Blebs were isolated from round cells synchronised in interphase or mitosis (as described in A) (Step 1). Blebbing was induced with treatment with the actin depolymerising drug Latrunculin B (Step 2). Blebs were detached from the cells with sheer stress (Step 3) and separated blebs were isolated from the cells using a 5 µm filter (Steps 4, 5). Re-assembly of a dynamic actin cortex was induced in blebs through addition of ATP regenerating system (see Methods) and the alpha-toxin Haemolysin, which permeabilises the blebs, allowing for ATP regenerating system uptake (Step 6). (E) Representative confocal images of the actin cortex in blebs isolated from interphase (top) and mitotic (bottom) cells. White: Phalloidin (F-actin) Insets: individual blebs. Scale bars= 20 µm; 0.5 µm.

We then purified blebs to enrich for actin cortex components from synchronised, rounded cells (Figure 1D) by adapting a previously published protocol (18) (see Methods for details). In brief, blebbing was induced using the actin depolymerizing drug Latrunculin B (20) and blebs were detached from cells by shear stress. We observed that mitotic HeLa cells yielded fewer blebs than the constitutively blebbing M2 cells used in previous studies (18, 19); to account for this limited output, we considerably increased the amounts of cells used. Furthermore, in our protocol blebs are isolated from non-adherent cells, we thus added a stringent filtration step to ensure the removal of entire cells from the cortex preparation (Figure 1D). As previously described (18), to facilitate cortex re-assembly, the bleb membrane was then permeabilised by addition of Hemolysin A and of anexogeneous ATP regeneration system (Figure 1D). This protocol allowed for the isolation of blebs from synchronised cells with a clearly defined actin cortex (Figure 1E). Bleb lysates were enriched in actin and actin-binding proteins and displayed lower levels of nuclear proteins compared to whole cell lysates (Supplementary Figure 1). Taken together, our purification protocol successfully isolated cortical fractions from cells synchronised in interphase and mitosis.

### Proteomic analysis of actin cortex-enriched blebs from interphase and mitotic cells

To identify proteins potentially involved in the regulation of cortical tension, we analysed blebs isolated from interphase and mitotic cells using liquid chromatography (LC)-tandem mass spectrometry (LC-MS/MS). For this, purified blebs from three biological replicates for each condition were lysed in denaturing Laemmli buffer and proteins were resolved by SDS-PAGE (Figure 2A). Proteins were then subjected to in-gel trypsin digestion, and purified peptides were analysed by LC-MS/MS. Overall, we identified 117,088 unique peptides from 2,268 unique proteins in interphase and mitosis (Figure 2B and C, Supplementary Table 1). For further analysis, we only considered proteins that were detected from at least 2 unique peptides and with an average spectral count of 2 or higher, in triplicates from both phases of the cell cycle, narrowing down the list to 922 proteins (Figure 2D, Supplementary Table 2). These criteria allowed us to focus on proteins that were consistently and reliably detected in our samples. Furthermore, focusing on proteins present in both interphase and mitosis eliminated multiple nuclear proteins that were detected at high levels in mitotic blebs but absent in interphase blebs (Supplementary Table 1), likely due to increased cytoplasmic levels of nuclear components following nuclear envelope breakdown. Out of the 922 proteins selected, myosin heavy chain IIA (MYH9), actin (ACTG1) and filamin A (FLNA) were the most abundant based on spectral counts, which reflect both protein abundance and size (Figure 2E). Overall, many of the most abundant proteins identified were actinrelated proteins (Figure 2E, magenta dots).

**Table 1.**
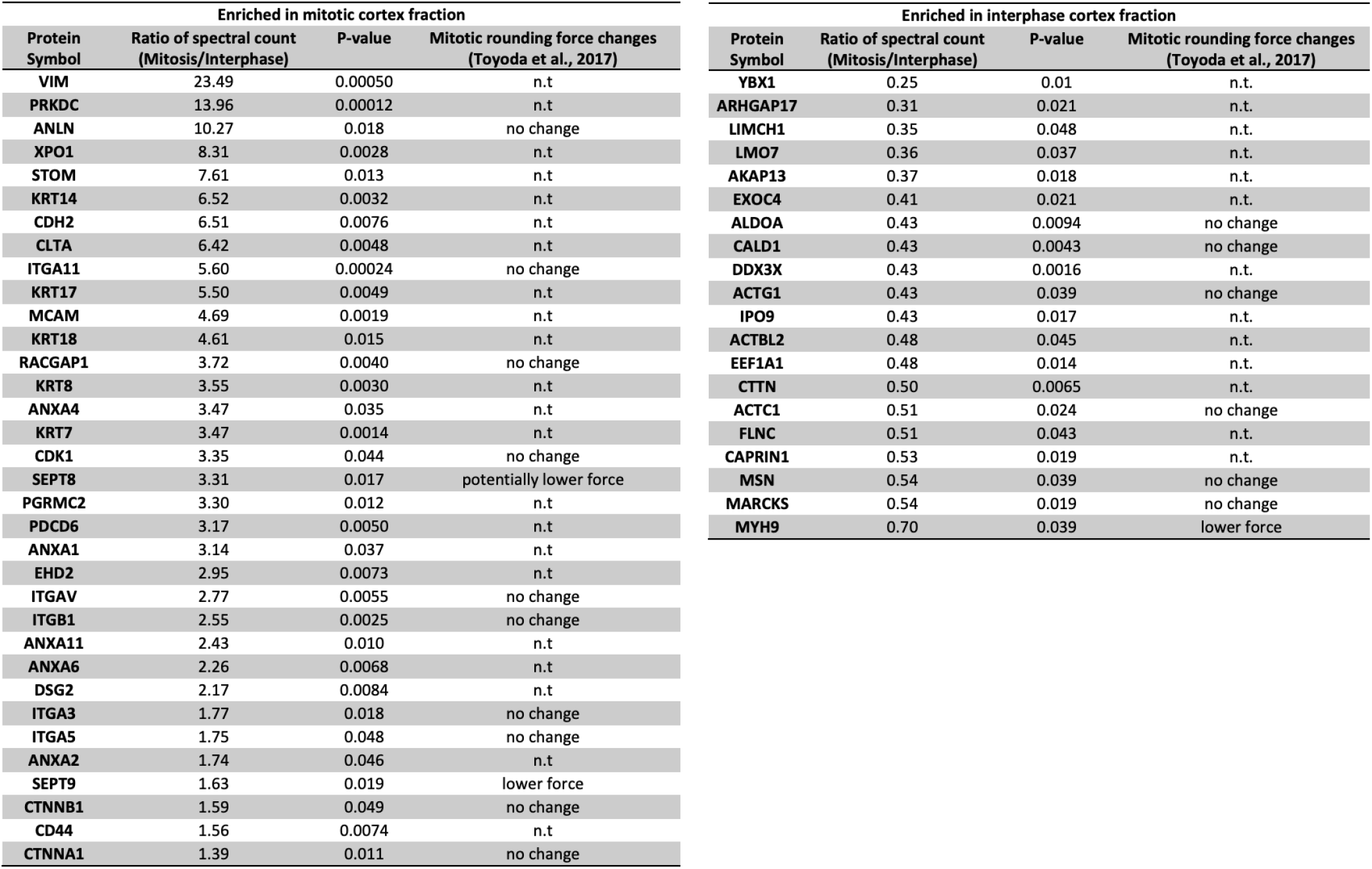
Actin-related proteins detected in isolated blebs showing a significant difference in levels between interphase and mitosis (P-value<0.05). Fold-change and exact P-values were calculated from normalised spectral counts detected in blebs in 3 experimental replicates. Right column: rounding force changes upon protein depletion, as reported in (24); n.t. (not tested): protein not examined in the mechanical screen in (24); no change: protein for which no change was detected with any of the esiRNA tested; potentially lower force: protein for which the change in rounding force was detected with one of the esiRNA sequences tested; lower force: protein for which rounding force changed with all esiRNA sequences tested (24).

**Fig. 2.**
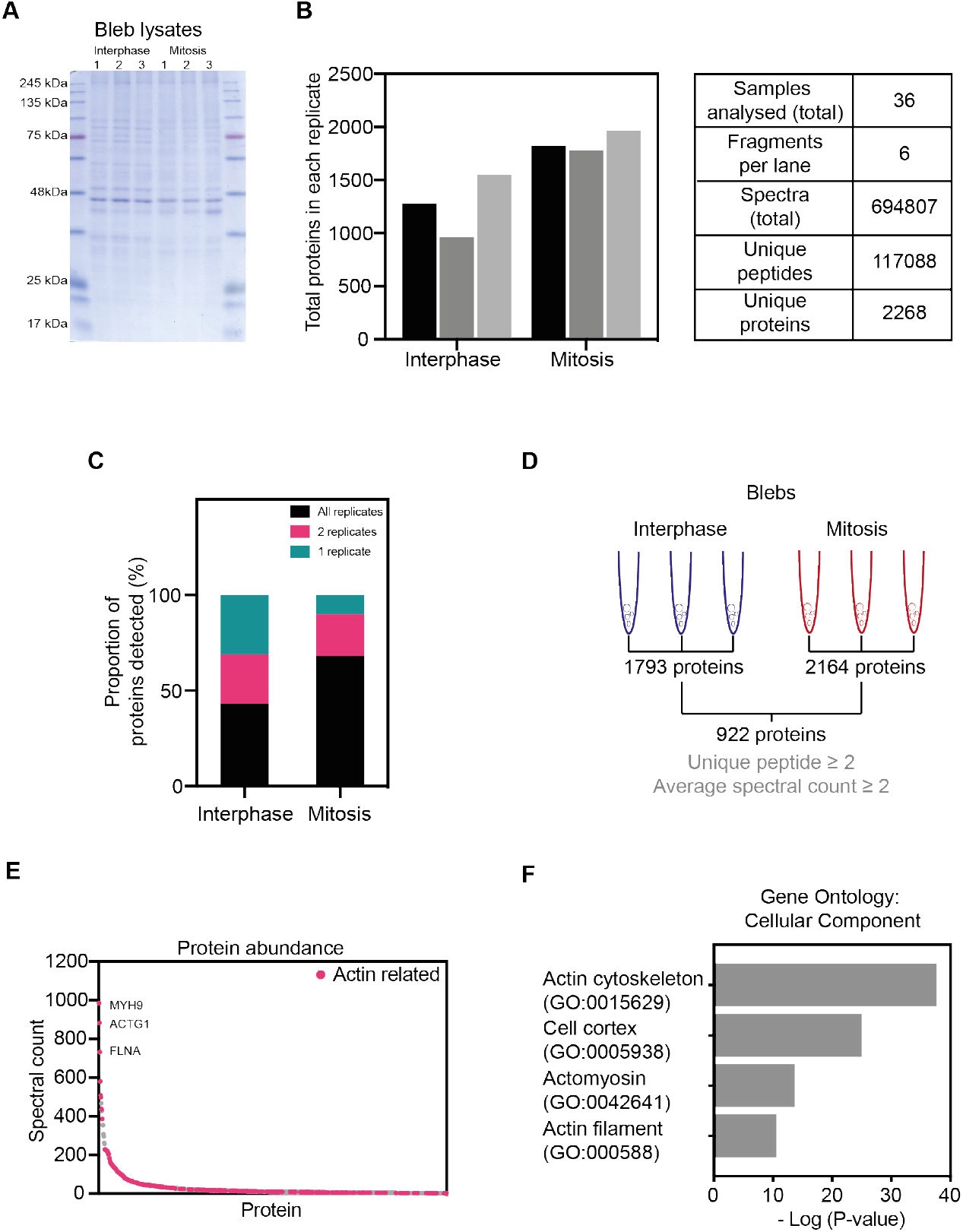
Proteomic analysis of blebs isolated from interphase and mitotic cells. (A) Coomassie staining of isolated blebs from interphase and mitosis in the three experimental replicates used for mass spectrometry. (B) Quantification of the number of proteins (left) and other overall readouts of the mass spectrometry analysis of samples from (A). (C) Percentage of proteins detected in all, two or one of the replicates. (D) Schematic summarising the process of protein selection for further analysis. Mass spectrometry detected 1793 and 2164 different proteins in blebs isolated from interphase and mitotic cells, respectively. Out of these, 922 proteins with a unique peptide number above 2 and normalised spectral count above 2 were present in both interphase and mitotic blebs; these proteins were selected for further analysis. (E) Average spectral counts in all replicates from interphase and mitotic blebs of the 922 proteins selected as described in (D). Magenta dots: actin related proteins. (F) Gene Ontology (GO) analysis of the selected 922 proteins, focusing on GO terms for cellular components related to the actin cortex.

To globally characterise the composition of the identified protein dataset, we next performed a Gene Ontology (GO) analysis for cellular components, molecular functions, and biological processes (Figure 2F, Supplementary Figure 2). Cellular component GO terms describing actomyosin (P-value= 3.78 × 10^*−*14^), actin filament, (P-value= 4.68 × 10^*−*11^), cell cortex (P-value= 1.60 × 10^*−*25^) and actin cytoskeleton (P-value= 3.74 × 10^*−*38^) were enriched for in our dataset (Figure 2F). Analysis of molecular functions, and biological processes GO terms further supports a high representation of actin-related proteins in isolated blebs (Supplementary Figure 2). Taken together, our observations suggest that isolated blebs are indeed enriched in actin cortex components and are thus a good model system to compare cortex composition between interphase and mitosis.

### Analysis of actin-related proteins in cortex-enriched blebs from interphase and mitotic cells

To identify potential regulators of cortical tension, we next compared the levels of individual proteins detected in blebs isolated from cells in interphase and mitosis (Figure 3A, Supplementary Table 2). Only 180 out of the 922 proteins detected displayed a significant difference in protein levels between interphase and mitosis (corresponding to a P-value = 0.05, data points above the dashed line in Figure 3A), suggesting that cortex composition is largely similar between these two phases of the cell cycle. To narrow down our candidate list, we focused on actin-related proteins. These were selected based on a previously published list of F-actin binding proteins identified by pull down of F-actin binding cellular fractions (16), complemented by manual curation to select further actin-related factors. Based on these criteria, we found 238 actin-related proteins in the cortex-enriched blebs (Figure 3A and B, Supplementary Table 1). This narrowed down list included many proteins known to directly bind and regulate actin filaments, as well as membrane and adhesion proteins, the intermediate filaments vimentin and keratin, and various Rho-GTPases and their regulators. Notably, the other proteins found in blebs included tubulins and microtubule-binding proteins, as well as multiple factors involved in intracellular trafficking, which might also indirectly interact with the actin cortex (Supplementary Tables 1, 2). Out of the 238 identified actin-related proteins, 54 significantly changed in levels between interphase and mitotic blebs to a P-value <0.05. While this cut-off is somewhat arbitrary given the small number of experimental replicates inherent to a mass spectrometry study, this reduced list represents actin-related proteins that most consistently changed levels in cortex-enriched blebs between interphase and mitotic cells (Table 1, Supplementary Figure 3).

**Fig. 3.**
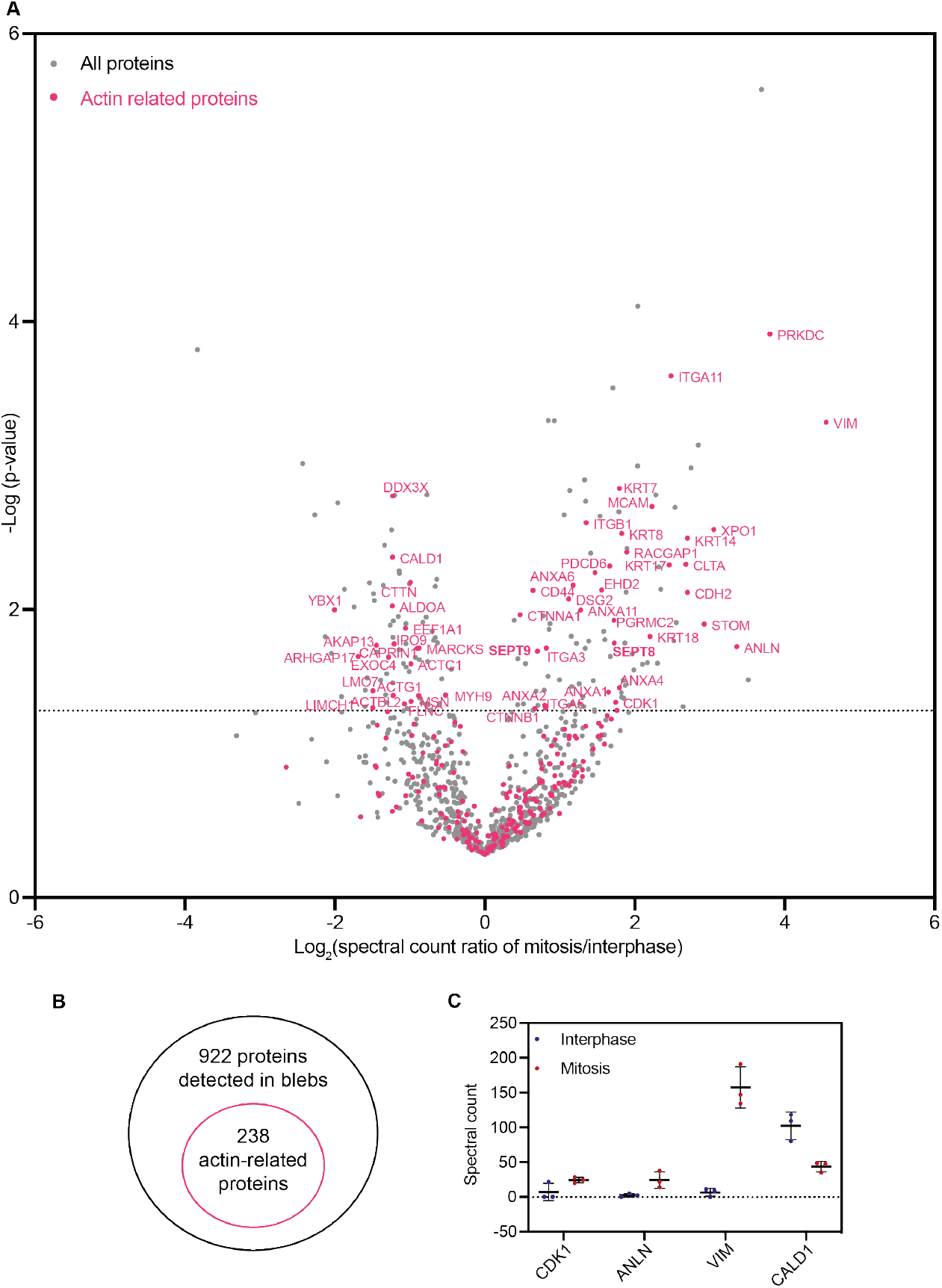
Changes in cortical levels of actin-binding proteins between interphase and mitosis. (A) Volcano plot of the 922 selected proteins detected in blebs, showing the enrichment (x axis) and the significance of this enrichment (p-values, y axis) between interphase and mitosis. Dotted line highlights -Log10 (P-value) = 1.3, corresponding to P-value= 0.05. Statistics: Student’s t-test. Magenta dots: actin related proteins. (B) Schematic of actin-related proteins among all proteins detected in blebs. (C) Spectral counts in the mass spectrometry analysis of mitotic and interphase blebs for proteins known to change in levels at the cortex between interphase and mitosis. Each data point corresponds to an individual replicate, with mean ± 1 standard deviation shown.

Several of the identified proteins had previously been shown to change levels between interphase and mitosis (Figure 3C). In particular, we found that the levels of cyclin-dependent kinase 1 (CDK1), a known cell-cycle regulator that increases in mitosis (21), and anillin, a cortical protein that translocates from the nucleus to the actin cortex in mitosis (22) were higher in mitotic compared to interphase blebs. Furthermore, the intermediate filament protein vimentin displayed strongly increased levels in mitotic blebs, consistent with recent findings demonstrating a role for vimentin in the mitotic cortex (15, 16). Finally, caldesmon displayed lower levels in mitotic compared to interphase blebs, consistent with previous observations showing that caldesmon dissociates from actin filaments during mitosis (23). Together, these observations suggest that our mass spectrometry analysis successfully identifies proteins for which cortical levels change between interphase and mitosis.

We then further scrutinised our dataset by comparing our list of 238 actin-related cortical proteins to a published targeted screen of proteins involved in the generation of the mitotic rounding force (24). We found that 103 out of the 238 actin-related proteins detected in blebs had been tested in this mechanical screen. Out of these, 10 were shown to reproducibly and significantly reduce mitotic rounding force upon depletion (Supplementary Table 3). These 10 regulators of cortex mechanics included the heavy chain of non-muscle myosin IIA (MYH9), which served as a positive control in the mechanical screen (24), upstream regulators of actomyosin dynamics (RAC1, ROCK), the septin SEPT9, and proteins involved in the control of actin organisation. This last category included proteins affecting actin polymerisation and nucleation (DIAPH1, PFN1, DBN1, CYFIP1), and actin bundling and crosslinking (ACTN4, FSCN1). Mechanisms through which actin length regulators (e.g. DIAPH1, PFN1) and crosslinkers (e.g. ACTN4) affect force generation in actomyosin networks have been explored previously (12, 25, 26). Furthermore, MYH9 and SEPT9 were the only two proteins out of the 10 validated cortex mechanics regulators, to also show a significant change in cortical levels in our mass spectrometry analysis (Table 1). We thus decided to focus on the role of septins in cortex regulation for the rest of the study.

### A role for the septin protein family in the regulation of the mitotic cortex

We then asked whether septins could play a role in the regulation of the cortex reorganisation between interphase and mitosis. Septins 2 and 9 have been previously shown to localise to the prometaphase cell cortex (27), but their role at the mitotic cortex remains unclear. Interestingly, our data indicate that the levels of several members of the septin family increase at the cortex between interphase and mitosis, with septin 8 and 9 displaying the most significant increase (Figure 3A, 4A, Supplementary Figure 4A). Proteins of the septin family assemble into higher order structures, including filaments, and are increasingly considered a component of the cytoskeleton (28, 29). Septins are generally classified into four different subgroups (SEPT3, SEPT2, SEPT7, SEPT6), with members of the same subgroups displaying some redundancy for the formation of oligomers and filaments. Septin 9 is the only member of SEPT3 subgroup that was detected in blebs, it also displayed a statistically significant change in cortical levels between interphase and mitosis in our analysis (Figure 3A and 4A, Supplementary Figure 3, Table 1), and had been shown to significantly affect mitotic rounding forces upon depletion in a mechanical screen (24). We thus focused on septin 9 for further investigation.

**Fig. 4.**
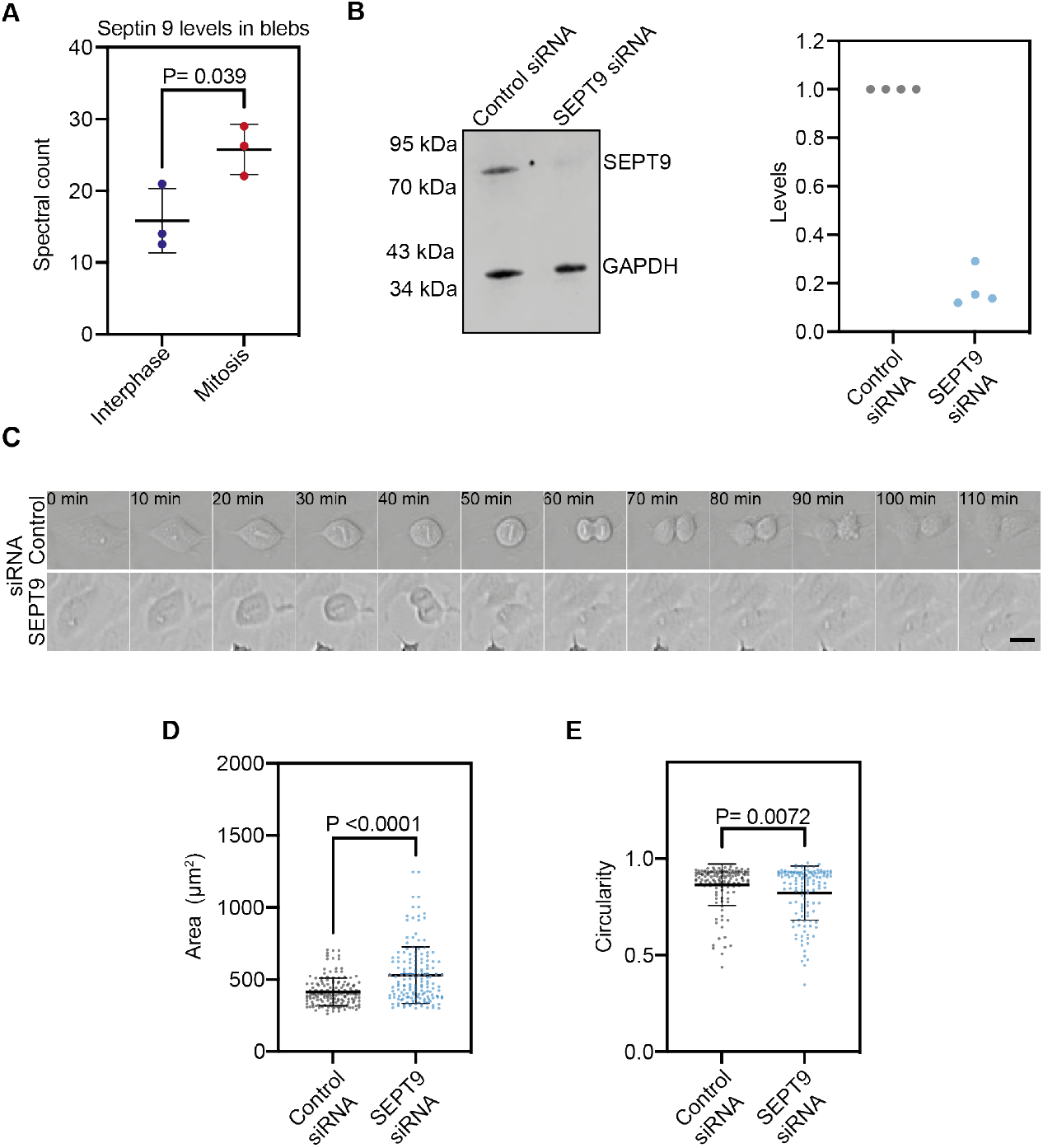
Septin 9 regulates mitotic rounding. (A) Spectral counts for septin 9 in the mass spectrometry analysis of mitotic and interphase blebs; each data point corresponds to an individual replicate, with mean ± 1 standard deviation shown (B) Example Western blot and quantification showing the decrease in septin 9 levels upon siRNA treatment. Membrane is representative of n= 4 samples used for quantification. Quantified levels were normalised to the loading control (GAPDH) and control siRNA conditions. (C) Timelapse (every 10 min) of cell division of cells treated with siRNA against septin 9 (bottom) and control siRNA (top). Scale bar= 20 µm. Quantification of cell area (D) and circularity (defined as 4*π*\*area/perimeter^2^, E) in mitosis of control and SEPT9 siRNA treated cells. Graph, mean ± 1 standard deviation, 3 independent experiments. Statistics: Unpaired t-test.

We first assessed whole cell levels of septin 9 and found that not only cortical (Figure 4A) but overall septin 9 levels increased between interphase and mitosis (Supplementary Figure 4B), suggesting that septin 9 might play an important role in cell division. We thus depleted septin 9 (Figure 4B and C) and examined the resulting effects on the cortex-driven mitotic rounding by analysing cell shape using live-cell imaging (Figure 4C-E, Supplementary Figure 4C, D, Supplementary Movies 1, 2). We found that septin 9-depleted cells showed strong mitotic rounding defects (Figure 4D and E, Supplementary Figure 4C and D). In particular, septin 9-depleted cells displayed larger and less circular cross-sections (Figure 4C-E), as well as a significantly higher aspect ratio (Supplementary Figure 4D) compared to control cells. Taken together, our dataset identifies septin 9 as an important regulator of cellular shape at the onset of mitosis, highlighting the effectiveness of our dataset for identification of cell surface mechanics regulators.

## Discussion

Here, we compared the composition of the actin cortex between interphase and mitosis, through mass spectrometry of cellular blebs isolated from synchronised rounded interphase and mitotic cells. Unlike previous studies, where proteic composition of whole cell lysates (30) or re-polymerised Factin fractions (16) were compared between interphase and mitosis, our approach directly assesses cellular fractions enriched for the actin cortex. As such, our study can detect differences in protein levels at the cortex due to changes in either protein expression or localisation. Out of the 922 proteins detected in both interphase and mitotic blebs, 238 were actin-related. These 238 proteins represent a candidate list for the regulation of the cortex remodelling upon mitosis entry (Supplementary Table 3).

Analysis of our candidate list pointed to the septin family as a potential and underexplored regulator of the mitotic cortex. Septins were first identified as key regulators of yeast cytokinesis (31). In mammalian cells, several septins have also been shown to interact with actomyosin networks and with the plasma membrane (32). Depletion of septins 2, 7, and 11 has been shown to affect cleavage furrow ingression, leading to multinucleated cells (27). In particular, septin 2 has been shown to colocalise with non-muscle myosin II in dividing cells, potentially acting as a scaffold to promote myosin phosphorylation during cytokinesis (33). Furthermore, septin 9 depletion has been shown to interfere with midbody abscission following cytokinesis (27). Whether septins also affect the cortex and cellular shape during earlier stages of mammalian cell division has received little attention. Our study suggests that septins, and septin 9 in particular, regulate the cortex already in the very early stages of mitosis, and contribute to the changes in cell surface mechanics that drive mitotic rounding. How exactly septin 9 is recruited to the mitotic cortex and how it regulates mitotic cortex mechanics and resulting cell rounding constitute interesting avenues for future studies.

The identification of septin 9 as a regulator of the cortex-driven cell shape changes at mitosis entry highlights the potential of the dataset we generated in detecting important cortex regulators amongst the multiple cortical components. More broadly, by systematically comparing the composition of the mitotic and interphase actin cortex, our study will be an important resource for investigations of mitotic shape changes, and of actin cortex mechanics in general.

## Methods

### Cell culture, cell synchronization and siRNA treatment

HeLa cells from MPI-CBG Technology Development Studio (TDS) were cultured in DMEM Glutamax (Thermo Fischer Scientific) supplemented with 10% fetal bovine serum (FBS) (Thermo Fischer Scientific); 1% PenicillinStreptomycin (Thermo Fischer Scientific); 1% L-Glutamine (Thermo Fischer Scientific). For passaging, cells were detached from culturing flasks with Trypsin-EDTA (Thermo Fischer Scientific). Cells were regularly tested for mycoplasma. Cells were synchronised in interphase with 2 mM thymidine for approximately 22 h, and in prometaphase with 2 µM S-Trityl-L-cystine (STLC) for 16 h. Mitotic cells were further enriched for with mitotic shake-off. Thymidine and STLC were removed from the cells before the bleb isolation. For siRNA depletion, cells were transfected with ON-TARGET plus Human SEPT9 pool siRNA (Horizon Discoveries, 006373-00) and Non-targeting Control pool (Horizon Discoveries, 001810-10) using Lipofectamine™ RNAiMAX Transfection Reagent (Thermo Fischer Scientific) in antibiotic-free media. Cells were treated with siRNA 24h before experiments were performed.

### Bleb isolation

Blebs were isolated from either mitotic cells synchronised with STLC and detached with mitotic shakeoff or cells synchronised in interphase with thymidine and detached with trypsin. Trypsin was deactivated with cell culturing media, followed by centrifugation at 1000 rpm for 3 min and exchanging media for fresh culturing media. Blebbing was induced by addition of either 1.7 µM (interphase) or 2.4 µM (prometaphase) of Latrunculin B (Sigma Aldrich), immediately followed by shaking on a horizontal benchtop shaker for 15 min at room temperature to detach the blebs from the cells. Latrunculin was then washed out through centrifugation of isolated blebs (and cells) at 4410 g for 6 minutes and re-suspension in intracellular buffer (0.33% 3 M sodium chloride, 28% 1 M pH 7.2 L-glutamic acid, 1.4% 1 M magnesium sulphate, 13.3% 0.1 M calcium chloride, 20.4% 0.1 M pH 7.2 EGTA, 4% 1 mM pH 7.2 HEPES, 32.5% dH2O, potassium hydroxide to adjust the pH). To separate entire cells and isolated blebs, cells were firstly pelleted with a 4 min centrifugation at 100 g. The supernatant was then filtered with 5 µm Satorious Minisart filters (FIL6602, Minisart) to remove remaining cells and any larger debris. Collected blebs were then pelleted with a centrifugation at 16100 g for 5 min and incubated in a solution containing an exogeneous ATP regeneration system (energy mix) and alpha-toxin to permeabilise the bleb membrane (5% A-Hemolysin alpha-toxin (1 mg/mL; H9395, Sigma Aldrich), 2% energy mix (50 mg/mL UTP, 50 mg/mL ATP, 255 mg/mL creatine phosphate, 2% creatine kinase (10 mg/mL), in intracellular buffer)) for 10 min, followed by centrifugation at 16100 g for 5 min and blebs resuspension in 500 µL re-suspension buffer (50% intracellular buffer, 44% dH20, 1% energy mix, 5% creatin kinase) for 20 min. Purified blebs were then lysed directly in the Laemelli sample buffer and the lysates were prepared for mass spectrometry.

### Mass spectrometry

To obtain sufficient material for MS analysis, isolated blebs were prepared from 15 of T175 flasks of cells synchronised in interphase and 60 flasks of cells synchronized in mitosis, in three experimental replicates for each phase of the cell cycle. For MS analysis, Coomassie-stained gel bands were excised and subjected to in-gel trypsin digestion, as described previously (34). The resulting peptides were extracted and subjected to capillary LC-MS/MS using a high-resolution hybrid mass spectrometer LTQ-orbitrap XL (Thermo Fisher Scientific). Experiments were performed in triplicates. Database searches were performed against The Uniprot SwissProt Human database (containing 20 347 protein entries) using PEAK Studio (version 8.5) as search engine, with trypsin specificity and three missed cleavage sites allowed. Methionine oxidation, Lysine acetylation, Cysteine carbamidomethylation, Serine/Threonine/Tyrosine phosphorylation and asparagine/glutamine deamidation were set as variable modifications. The fragment mass tolerance was 0.01 Da and the mass window for the precursor was ± 10 ppm. The data were visualised with scaffold (version 4.8.6) and minimum number of peptides per protein was set to 2 for data analysis. Data availability: The raw mass spectrometry datasets have been deposited in ProteomeXchange through partner MassIVE as a complete submission and assigned MSV000088744 and PXD031308. The data can be downloaded from ftp://MSV000088744@massive.ucsd.edu using the password blebs.

### Gene Ontology and candidate list curation

For Gene Ontology (GO) analysis the statistical overrepresentation test function from PantherDB (http://www.pantherdb.org/) was used. Actin-related proteins from the bleb extracts were identified by comparing the list with previous mass spectrometry analysis of the F-actin interactome in interphase and mitotic cells (16), and by manually adding known actin related proteins.

### Western blots

Cells were lysed directly in Laemelli sample buffer, boiled and sonicated. 30 µg of total protein per sample was loaded on NuPage 4-12% Bis-Tris Protein gels (Thermo Fischer Scientific) or 4–15% Mini-PROTEAN TGX StainFree Protein gels (Biorad) and run at 200 V as per manufacturer’s instructions. Proteins were transferred to a nitrocellulose membrane (Thermo Fischer Scientific) using the BioRad transfer system at 100 V for 60 min at 4°C or with Trans-Blot Turbo Mini PVDF Transfer Packs using Trans-Blot Turbo Transfer System (Biorad). The membrane was blocked for 30 min with the Odyssey Blocking Buffer (TBS) (Licor) and stained overnight with primary antibodies at 4°C. Primary antibodies used: phospho-histone H3 (Cell Signalling, 9713S), septin 9 (Sigma-Aldrich, HPA042564) and GAPDH (Abcam, ab8245) in 5% milk in PBS with 0.1% tween, diluted 1:500, 1:1000 and 1:5000, respectively. Following primary antibody incubation, membranes were washed three times with PBS 0.1% tween. Licor secondary antibodies conjugated to IRDyes were diluted 1:5000 in 5% milk in PBS 0.1% Tween 20 and incubated with the membranes at room temperature for 60 min. Membranes were imaged with Odyssey FC system and the results were analysed with the Studio Lite software.

### Preparation of fixed samples for imaging

Samples, isolated blebs and cells were spun on poly-L-lysine coated 25 mm coverslips by centrifugation at 460 g for 10 min. Cells were fixed with 4% paraformaldehyde (PFA) in PBS for 10 min followed by 10 min permeabilisation with 0.2% Triton X-100 at room temperature. Blebs were fixed with combined permeabilisation-fixation for 6 min with 4% PFA in intracellular buffer with 0.2% Triton X-100 followed by 14 min fixation with 4% PFA in intracellular buffer at room temperature, followed by three washes with PBS. Samples were stained with DAPI and Phalloidin-Alexa568 (1:500 dilution from stock in MeOH) for 1 h, followed by three washes with PBS. Samples were imaged using Olympus FluoView FV1200 Confocal Laser Scanning Microscope using a 60x oil objective (NA 1.4).

### Live imaging and analysis

Cells we treated with siRNA for 24 h before the start of the overnight imaging. 30 min prior to imaging, the culture media was changed to Leibovitz’s L-15 (Thermo Fischer Scientific) media supplemented with 10% FBS and 1% Penicillin-Streptomycin. Cells were imaged for 15 h at 37°C with 2 min time steps on an inverted Olympus IX81 microscope, controlled via Velocity interface, and a Hamamatsu Flash 4.0v2 ScMOS camera. Imaging was performed with the brightfield setting under a 20x air objective. Cell shape was analysed with manual segmentation using the measure feature of the FIJI image analysis software (35).

### Statistical analysis

Mass spectrometry results were analysed using Excel Microsoft and Prism (GraphPad Software) was used for other experiments. To compare means Student’s t-test was used.

## ACKNOWLEDGEMENTS

We thank all present and past Paluch lab members for critical discussions, Kevin Chalut, Ruby Peters and Iskra Yankieva for comments on the manuscript, Guillaume Charras for feedback on the project and manuscript. Proteomic analyses were performed by the Center for Advanced Proteomics Analyses (CAPCA), a Node of the Canadian Genomic Innovation Network that is supported by the Canadian Government through Genome Canada. This work was supported by the Medical Research Council UK (MRC Programme Award MC_UU_12018/5 to EKP), the European Research Council (Consolidator Grant 820188-NanoMechShape to EKP), and grants from the Canadian Institutes for Health Research (MOP-142374 and PJT-152995 to P.P.R). P.P.R. is a Senior Scholar of the Fonds de la recherche du Québec—Santé (FRQS). N.V. was funded by a MRC LMCB PhD studentship.

## AUTHOR CONTRIBUTIONS

Authors contributions N.V., P.P.R., and E.K.P. designed the research; N.V. carried out most of the experiments. G.L. helped with mass spectrometry; N.V., S.N. and P.P.R. analysed the mass spectrometry data; N.V., P.P.R., E.K.P. wrote the paper; M.S. provided technical support and conceptual advice. Funding Acquisition, P.P.R., and E.K.P. All authors discussed the results and manuscript.

