## Supplementary Information for "Proteomic analysis of the actin cortex in interphase and mitosis"

### Supplementary Figures

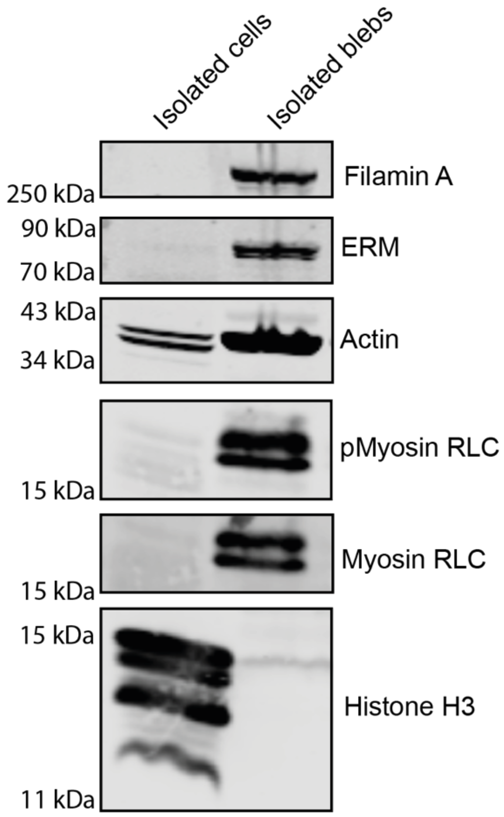


**Figure S1: Comparison of isolated blebs and cells.** Western blot for filamin A, ERM (ezrin, radixin, moesin), actin, myosin regulatory light chain (RLC), phospho myosin RLC, actin and nuclear protein (histone H3) in unsynchronised blebs and whole cells after an equal amount of total protein was loaded for a Western blot analysis.

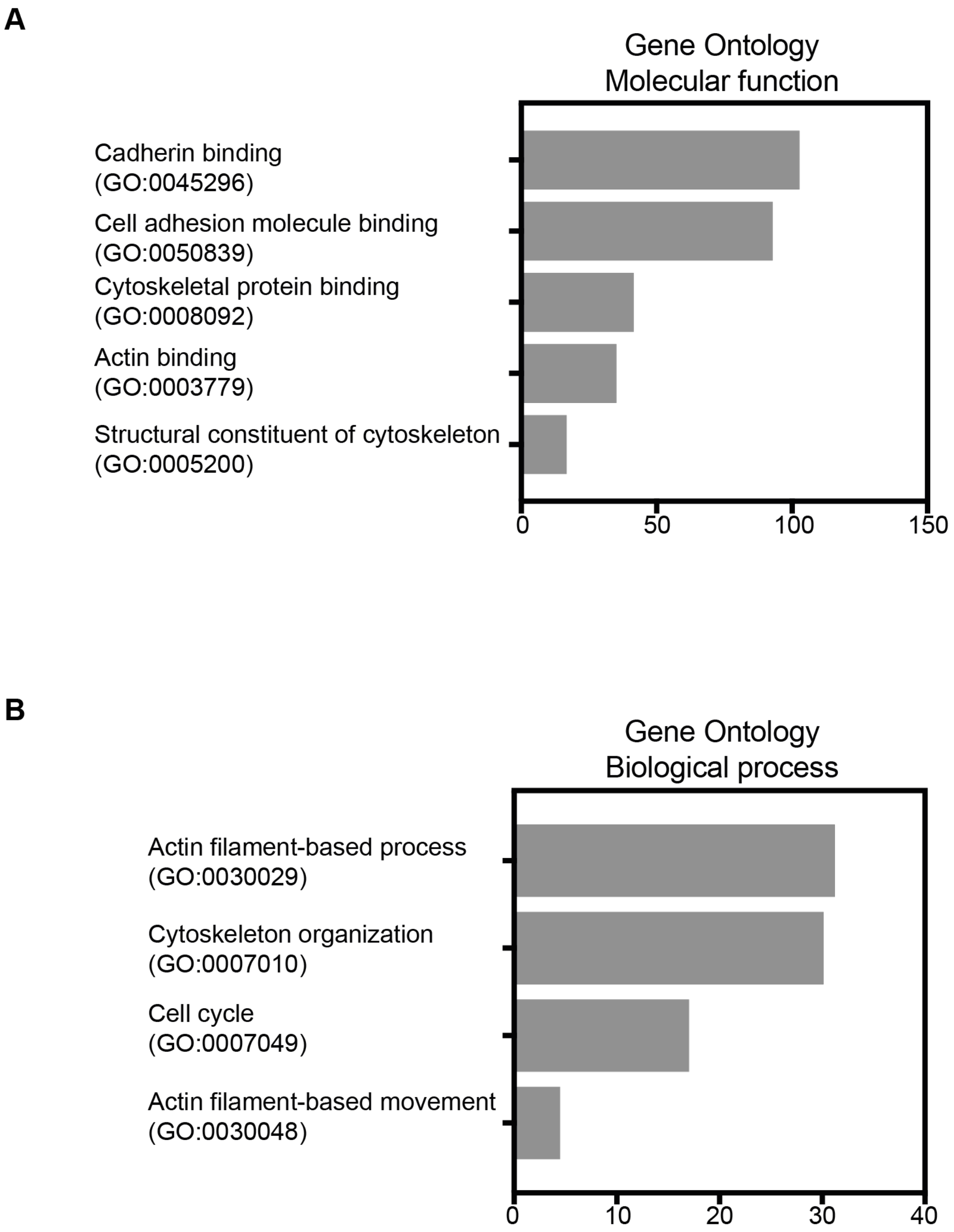


**Figure S2: Proteic composition of blebs.** Gene Ontology (GO) analysis of the selected 922 proteins, focusing on GO terms for molecular function (A) and biological function (B) related to the cell cortex.

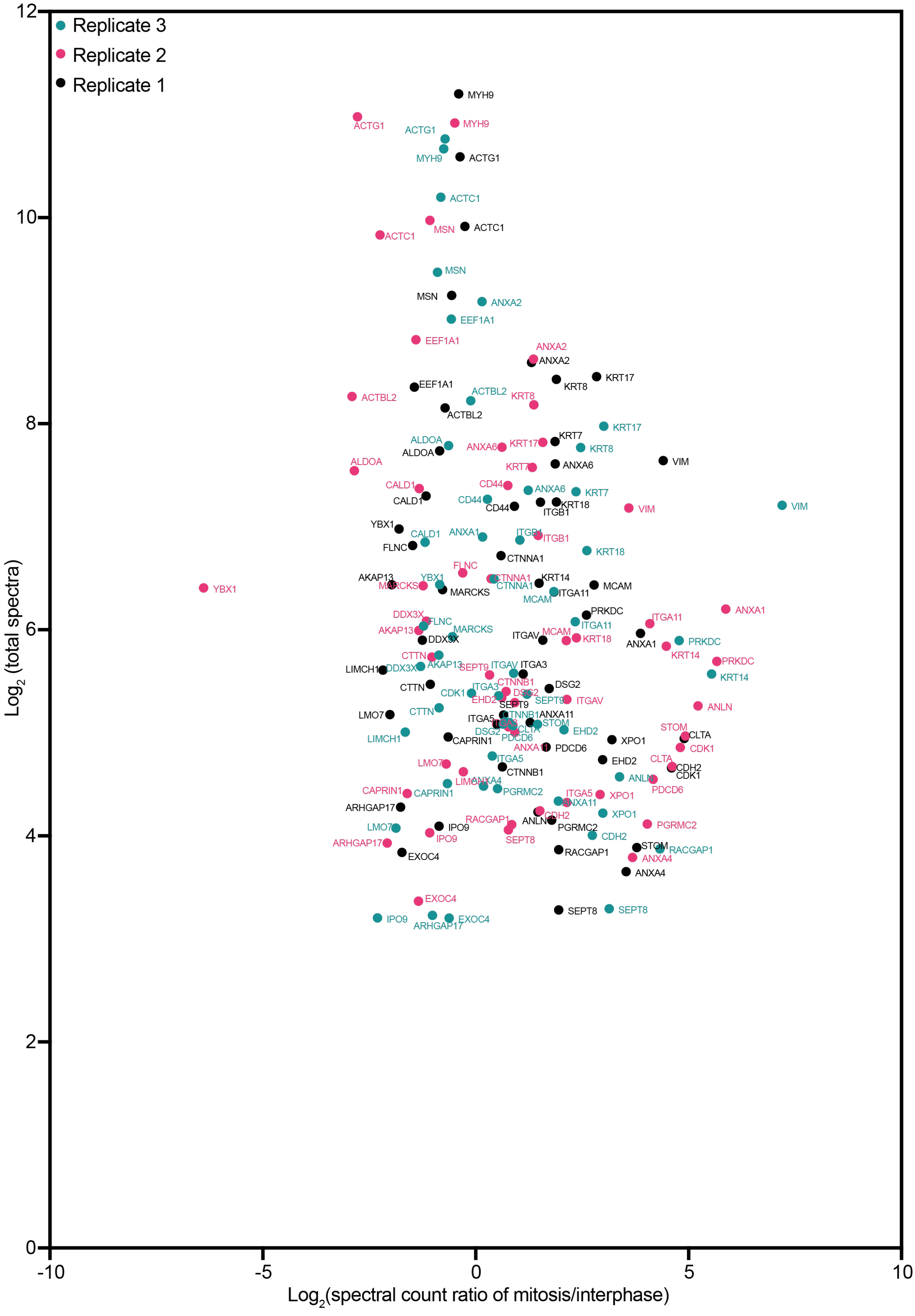

Replicate 3
Replicate 2
Replicate 1
12
10
8
6
4
2
0
-10
-5
0
5
10
Log2 (total spectra)
Log2(spectral count ratio of mitosis/interphase)
MYH9
ACTG1
ACTG1
MYH9
MYH9
ACTG1
ACTC1
MSN
ACTC1
ACTC1
MSN
MSN
ANXA2
EEF1A1
EEF1A1
ANXA2
ANXA2
KRT17
KRT8
EEF1A1
ACTBL2
KRT8
ACTBL2
ACTBL2
KRT17
ALDOA
ALDOA
ANXA6
KRT17
KRT7
KRT8
VIM
ALDOA
KRT7
ANXA6
CALD1
CD44
ANXA6
KRT7
CALD1
CD44
CD44
KRT18
ITGB1
VIM
VIM
YBX1
CALD1
ANXA1
ITGB1
ITGB1
FLNC
KRT18
CTNNA1
FLNC
AKAP13
YBX1
CTNNA1
KRT14
MARCKS
CTNNA1
MCAM
YBX1
MARCKS
ITGA11
MCAM
PRKDC
ANXA1
DDX3X
FLNC
ITGA11
ITGA11
AKAP13
MARCKS
DDX3X
ITGAV
MCAM
KRT18
ANXA1
PRKDC
KRT14
CTTN
AKAP13
SEPT9
ITGAV
ITGA3
PRKDC
LIMCH1
DDX3X
KRT14
ITGA3
CTNNB1
CTTN
CDK1
DSG2
DSG2
EHD2
SEPT9
ITGAV
SEPT9
ANLN
LMO7
CTTN
ITGA5
CTNNB1
ANXA11
STOM
DSG2
CLTA
EHD2
STOM
CLTA
LIMCH1
CAPRIN1
PDCD6
XPO1
ANXA11
PDCD6
CDK1
ITGA5
EHD2
CLTA
LMO7
CTNNB1
CDH2
CDK1
ANXA4
ANLN
PDCD6
CAPRIN1
CAPRIN1
PGRMC2
ITGA5
XPO1
ARHGAP17
ANXA11
XPO1
RACGAP1
ANLN
CDH2
LMO7
IPO9
PGRMC2
PGRMC2
IPO9
SEPT8
CDH2
ARHGAP17
STOM
RACGAP1
RACGAP1
ANXA4
EXOC4
ANXA4
EXOC4
SEPT8
SEPT8
IPO9
ARHGAP17
EXOC4

**Figure S3: Changes in levels of actin binding proteins in blebs between interphase and mitosis in experimental replicates.** (A) Volcano plot of 54 actin-related proteins that significantly change in levels in blebs from interphase compared and mitotic cells (listed in Table1), showing enrichment between interphase and mitosis (x-axis) and combined total of spectra (y-axis). Datapoints for all three experimental replicates are shown.

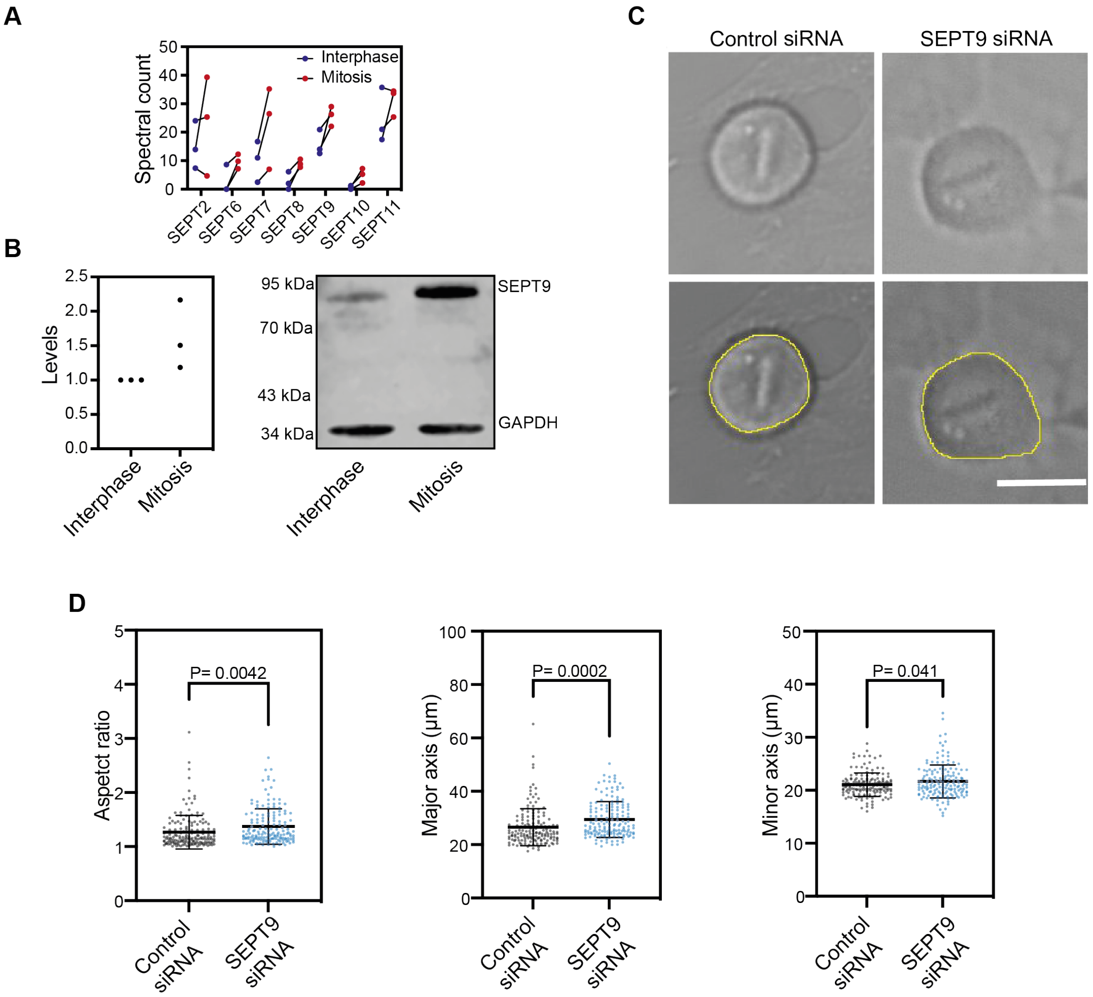


**Figure S4: Role of septins in mitosis.** (A) Levels of septins detected in blebs isolated from interphase and mitotic cells. Datapoints for all three experimental replicates are shown. (B) Western blot and quantification of septin 9 levels in mitotic and interphase whole cell lysates, normalised to loading control GAPDH relative to interphase levels. (C) Example of brightfield images of the cellular midplane of live mitotic cells treated with control and SEPT9 siRNA, that were used to analyse cell shape parameters (panel D and also Figure 4 D, E). Scale bar = 10 µm. (D) Quantification of cellular aspect ratio, major axis, and minor axis in mitosis in control and SEPT9 siRNA treated cells. Graph, mean ± 1 standard deviation, 3 independent experiments. Statistics: Unpaired t-test.

### Supplementary Tables

**Supplementary Table 1: Proteins detected in blebs.** Spectral counts of all proteins detected by mass spectrometry of isolated blebs, normalised to total protein levels in each replicate.

**Supplementary Table 2: Proteins detected in both interphase and mitotic blebs**. Mean spectral counts in interphase and mitosis, ratio of the means between mitosis and interphase (calculated from 3 replicates), and P-value calculated with Student's t-test, for the 922 proteins detected in both interphase and mitosis isolated bleb samples. Manually selected actin-related proteins (see Main text for list curation criteria) are highlighted in yellow.

**Supplementary Table 3**: **Actin-related proteins detected in blebs.** Average PAI (spectra count normalised to molecular weight), ratio of the means, and P-value (calculated with Student's t-test) for the 238 actin-related proteins detected with blebs. Right column: rounding force changes upon esiRNA treatment against the proteins (as reported in 24). Not tested: protein not examined; No change: protein for which no change was detected with any of the esiRNA tested; Potentially lower force: protein for which the change in rounding force was detected with some of the esiRNA sequences tested; Lower force: protein for which rounding force changed with all esiRNA sequences tested in (24).

### Supplementary Movies

**Supplementary Movie 1: Dividing cell transfected with control siRNA.** Movies from live-cell imaging were used for quantification of cell shape (Figure 4D-E, Supplementary Figure 4D). Time resolution= 2 min. Scale bar = 20 μm.

**Supplementary Movie 2: Dividing cell transfected with SEPT9 siRNA.** Movies from live-cell imaging were used for quantification of cell shape (Figure 4D-E, Supplementary Figure 4D). Time resolution= 2 min. Scale bar = 20 μm.
